# Oocytes can repair DNA damage during meiosis via a microtubule-dependent recruitment of CIP2A-MDC1-TOPBP1 complex from spindle pole to chromosomes

**DOI:** 10.1101/2022.11.04.514992

**Authors:** Jiyeon Leem, Jae-Sung Kim, Jeong Su Oh

## Abstract

Because DNA double-strand breaks (DSBs) greatly threaten genomic integrity, effective DNA damage sensing and repair are essential for cellular survival in all organisms. However, DSB repair mainly occurs during the interphase and is repressed during mitosis. Here, we show that, unlike mitotic cells, oocytes can repair DSBs during meiosis through microtubule-dependent chromosomal recruitment of the CIP2A-MDC1-TOPBP1 complex from spindle poles. After DSB induction, we observed spindle shrinkage and stabilization, as well as BRCA1 and 53BP1 recruitment to chromosomes and subsequent DSB repair during meiosis I. Moreover, p-MDC1 and p-TOPBP1 were recruited from spindle poles to chromosomes in a CIP2A-dependent manner. This pole-to-chromosome relocation of the CIP2A-MDC1-TOPBP1 complex was impaired not only by depolymerizing microtubules but also by depleting CENP-A or HEC1, indicating that the kinetochore/centromere serves as a structural hub for microtubule-dependent transport of the CIP2A-MDC1-TOPBP1 complex. Mechanistically, DSB-induced CIP2A-MDC1-TOPBP1 relocation is regulated by PLK1 but not by ATM activity. Our data provide new insights into the critical crosstalk between chromosomes and spindle microtubules in response to DNA damage to maintain genomic stability during oocyte meiosis.

## Introduction

DNA double-strand breaks (DSBs) greatly threaten the integrity of the eukaryotic genome, and unrepaired DSBs can compromise genomic integrity, causing developmental disorders, cell death, or cancer [1]. To counteract this, cells have evolved a variety of pathways to respond to DNA damage, collectively termed the “DNA damage response” (DDR). In response to DSBs, the master kinase ATM is initially activated and triggers a cascade of DDR events on chromatin-flanking DSBs [2]. The key event driving this process is the phosphorylation of histone H2AX at Ser139 (referred to as “γ-H2AX”), which serves as a docking platform for recruitment of the scaffolding protein MDC1 [3]. Once bound to γ-H2AX, MDC1 is phosphorylated by ATM and recruits multiple DDR factors, allowing further recruitment of ATM and the spread of γ-H2AX signaling to neighboring chromatins [4, 5]. Moreover, phosphorylated MDC1 interacts with RNF8, which in turn promotes RNF8/RNF168-dependent ubiquitination of γ-H2AX [6–9]. These signaling cascades further promote downstream repair factors, such as BRCA1 and 53BP1, to ensure DNA damage repair and checkpoint activation in response to DSBs [6–8, 10].

During the interphase, cells efficiently mount a coordinated response to DNA damage, activating cell cycle checkpoints and DNA repair pathways. However, during mitosis, cells are refractory to DNA damage and fail to mount DNA damage-induced cell cycle arrest [11–13]. During mitosis, the initial events, including H2AX phosphorylation and recruitment of MDC1, appear to be intact, but the recruitment of downstream DNA repair factors, such as RNF8, 53BP1, and BRCA1, is blocked [11, 14]. Instead, MDC1 recruits TOPBP1 to mitotic DSB sites, and TOPBP1 then forms a filamentous structure that bridges MDC1 foci and tethers broken chromosome ends [15]. Therefore, mitotic cells can efficiently mark DSB sites for repair in the subsequent G1 phase. The forced activation of DNA repair during mitosis causes sister telomere fusions, leading to missegregation of whole chromosomes [16]. Therefore, it is likely that normal cells have evolved to silence DNA repair mechanisms downstream of DNA damage robustly during mitosis to prevent the repair of broken DNA ends and deprotected telomeres, which would lead to mitotic catastrophe.

Unlike mitosis, meiosis consists of two consecutive cell divisions without an intervening S-phase, which is essential for reducing the ploidy. In the first meiotic division (meiosis I), homologous chromosomes pair and then segregate from each other, whereas sister chromatids segregate during the second meiotic division (meiosis II). A unique characteristic of meiosis in females, not seen in any other cell type, is prolonged arrest at the prophase of meiosis I, which is characterized by the presence of the germinal vesicle (GV). This extended arrest makes oocytes acutely susceptible to the accumulation of DNA damage [17]. Because prophase arrest in oocytes is equivalent to G2 phase in mitotic cells, one would expect oocytes to employ DDR mechanisms similar to that in mitotic cells. However, recent studies have revealed that oocytes do not induce a robust G2/M checkpoint in response to DNA damage [18, 19]. Therefore, oocytes with DNA damage resume meiosis and undergo GV breakdown (GVBD), unless the damage is not substantial. However, oocytes with DNA damage halted meiotic progression at the metaphase of the first meiosis (MI) by activating the spindle assembly checkpoint (SAC) [20, 21]. This DNA damage-induced SAC arrest is neither dependent on ATM/ATR nor associated with aberrant kinetochore-microtubule (kMT) attachments [22]. Moreover, there are strong indications that DNA repair occurs in oocytes during meiosis, unlike in mitotic cells, implying that the DDR in oocytes differs from that in somatic cells. Indeed, recent studies have revealed that DSB repair occurs in oocytes during GV or MII arrest, indicating that oocytes are equipped with DDR machinery and have the capacity to repair damaged DNA [20, 23, 24].

Despite recent advances in understanding DDR mechanisms in oocytes, it remains unclear how oocytes deal with DNA damage during meiosis I. Therefore, in the present study, we investigated whether oocytes could repair DSB during meiosis I. We found that oocytes can repair DSBs during meiosis I through microtubule-dependent chromosomal recruitment of the CIP2A-MDC1-TOPBP1 complex from the spindle pole via the kinetochore/centromere.

## Results

### Oocytes can repair DSBs by recruiting BRCA1 and 53BP1 during meiosis I

DNA repair pathways are suppressed during mitosis because the downstream repair factors BRCA1 and 53BP1 do not localize to mitotic chromosomes [11, 14]. We attempted to determine whether this is the case for oocytes during meiosis. To this end, oocytes at metaphase I (MI) stage were treated with etoposide (ETP) for 30 min, and the recruitment of BRCA1 and 53BP1 on metaphase chromosomes was examined. We found that BRCA1 and 53BP1 signals, which were barely detectable in control oocytes, significantly increased on chromosomes with strong enrichment at the centromeres after ETP treatment and returned to basal levels after 2 h of recovery (Figs. 1A–1C). Similarly, TUNEL signals increased after ETP treatment and decreased after 2 h of recovery (Figs. 1D and 1E). These results suggest that, unlike mitotic cells, oocytes can repair DNA damage during meiosis by recruiting the downstream repair factors BRCA1 and 53BP1 (Fig. 1F).

**Fig. 1.**
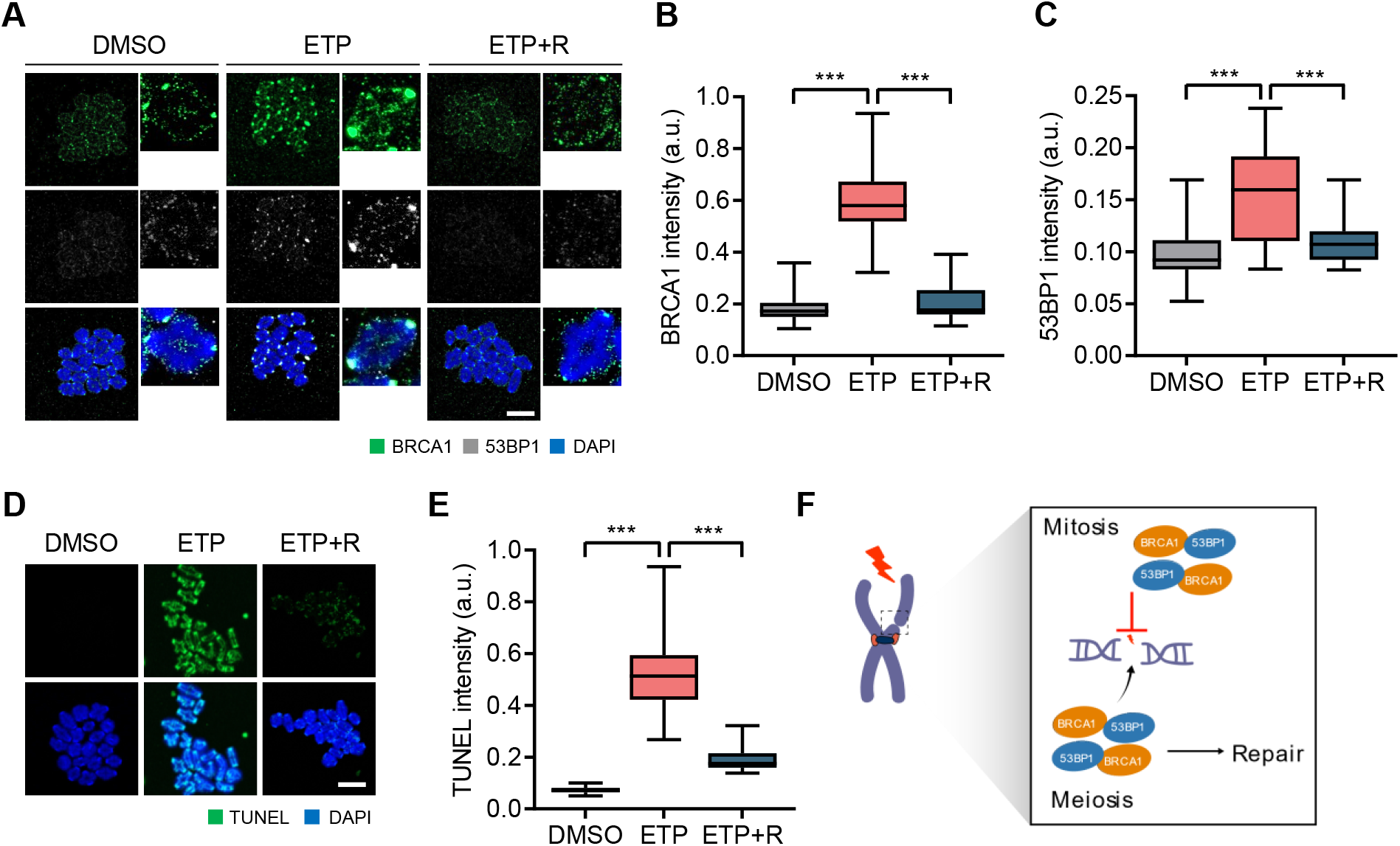
DSB repair and recruitment of BRCA1 and 53BP1 to chromosomes during meiosis I in oocytes. (A–E) Oocytes at the MI stage were treated with etoposide (ETP) for 30 min and recovered from DNA damage for 2 h (ETP+R). Control oocytes were treated with DMSO. (A) Representative images of metaphase chromosome spreads stained with BRCA1 and 53BP1 antibodies. Scale bar, 10 μm. (B, C) Quantification of BRCA1 and 53BP1 intensities. Data are presented as the mean ± SEM from three independent experiments. ***p < 0.0001. (D) Representative images of chromosome spreads showing TUNEL signals. Scale bar, 10 μm. (E) Quantification of TUNEL intensity. Data are presented as the mean ± SEM from three independent experiments. ***p < 0.0001. (F) Schematic diagram depicting the difference of DSB repair between mitotic cells and oocytes.

### DSBs induce chromosomal recruitment of MDC1 and spindle shrinkage and stabilization

To gain mechanistic insights into DSB repair during oocyte meiosis, we examined the initial events that occur after DSB induction. We found that γ-H2AX signals were detectable in intact oocytes and did not increase after ETP treatment, which is consistent with previous reports that metaphase chromosomes of MI oocytes are γ-H2AX positive regardless of DNA damage [25] (Figs. 2A and 2B). However, MDC1 signals were barely detectable but appeared on the chromosomes after ETP treatment (Fig. 2C). Interestingly, p-MDC1 signals were observed at the spindle poles and kinetochores in MI oocytes (Fig. 2C; Fig. S1), consistent with a previous report that MDC1 regulates CEP192-mediated microtubule-organizing center (MTOC) assembly during oocyte meiosis [26]. However, after ETP treatment, p-MDC1 signals were enriched in chromosomes (Fig. 2C; Fig. S1), suggesting that MDC1 can be a reliable DSB marker instead of γ-H2AX in MI oocytes and that MDC1 changes its localization in response to DNA damage.

**Fig. 2.**
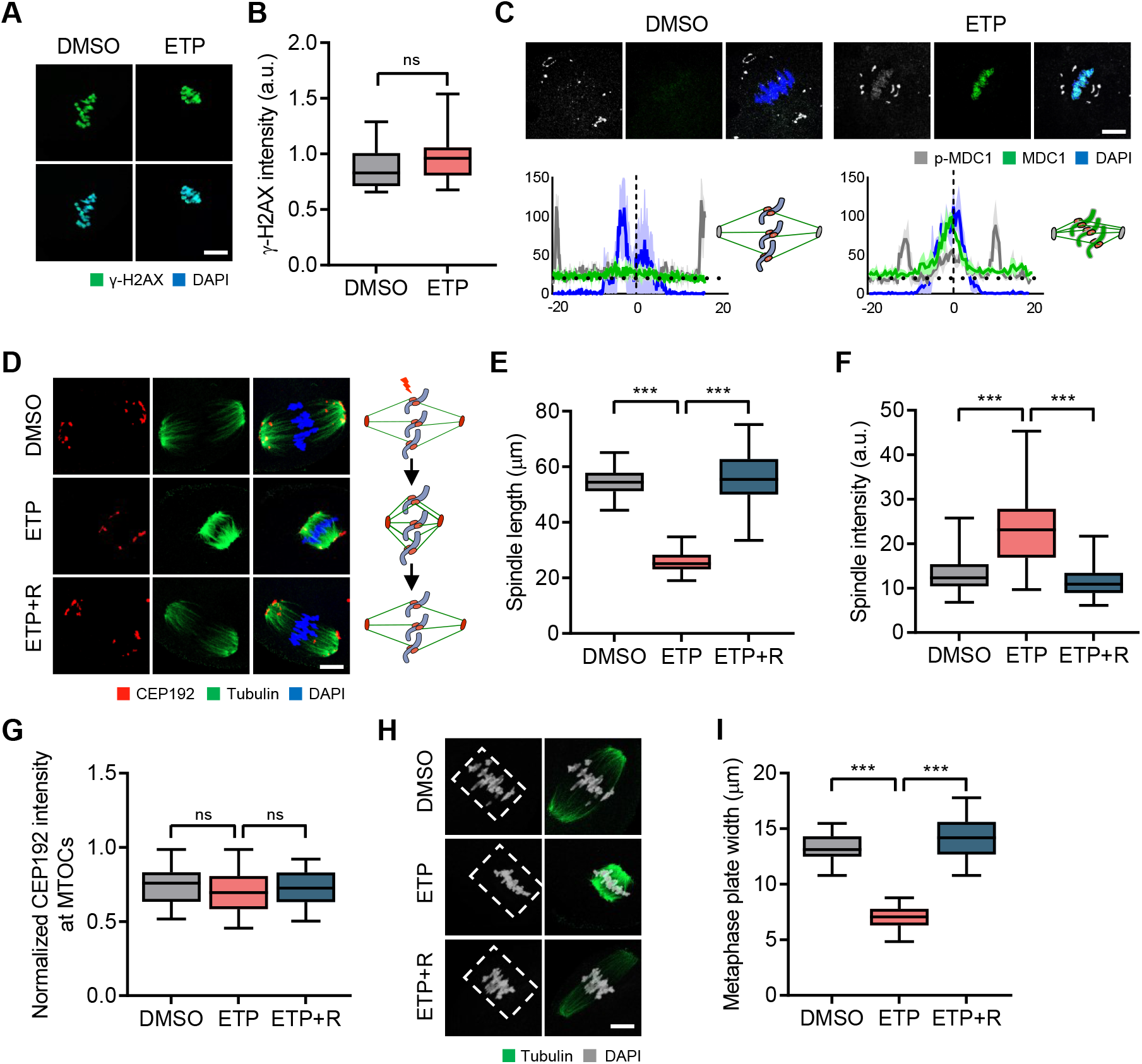
Chromosomal recruitment of p-MDC1 and dynamic changes in spindle and chromosome organization following DSB induction. (A) Representative images of control and ETP-treated MI oocytes stained with γ-H2AX antibody. Scale bar, 10 μm. (B) Quantification of γ-H2AX intensity. Data are presented as the mean ± SEM from three independent experiments. ns, not significant. (C) Representative images of MI oocytes stained with MDC1 and p-MDC1 antibodies. Scale bar, 10 μm. Profile of MDC1 and p-MDC1 signals between opposite spindle poles is shown with schematic illustration of spindle and chromosomes. (D) Representative images and schematic illustration showing spindle microtubules and MTOCs after ETP treatment and recovery (ETP+R). Scale bar, 10 μm. (E–G) Quantification of spindle length and intensity and CEP192 intensity at MTOCs. Data are presented as the mean ± SEM from three independent experiments. ***p < 0.0001. ns, not significant. (H) Representative images of spindle and chromosomes in MI oocytes. Scale bar, 10 μm. (I) Quantification of metaphase plate width. ***p < 0.0001.

In addition to the chromosomal relocation of p-MDC1, we noticed that p-MDC1 signals at the spindle poles appeared to be closer to the chromosomes after ETP treatment (Fig. 2C). This observation led us to investigate the impact of DNA damage on meiotic spindle organization. Notably, we found that ETP treatment markedly reduced the spindle length but did not impair kMT attachment (Figs. 2D and E; Figs. S2A and S2B). Moreover, the overall spindle and k-fiber intensities increased after ETP treatment (Figs. 2D and 2F; Fig. S2C). However, the intensity of CEP192 did not change after ETP treatment (Figs. 2D and 2G), suggesting that spindle pole integrity was not affected by DNA damage. In addition to spindle changes, chromosomes were more tightly aligned in the middle of the spindle, decreasing the metaphase plate width (Figs. 2H and 2I). This DSB-induced change in spindle and chromosome organization was rescued by 2 h of recovery (Fig. 2D–2I; Fig. S2). Therefore, our results suggest that DSBs induce spindle shrinkage and stabilization, as well as p-MDC1 relocation, implying crosstalk between meiotic bivalent chromosomes and spindle microtubules in response to DNA damage.

### Spindle microtubules are required to repair DSBs during oocyte meiosis

Because p-MDC1 is mainly localized at the spindle poles but is recruited to chromosomes with accompanying spindle shrinkage after ETP treatment, we hypothesized that spindle microtubules may play a role in the DDR during oocyte meiosis. To test this, MI oocytes were treated with ETP and allowed to recover from DNA damage in the presence of nocodazole. We found that nocodazole treatment impaired the ETP-induced chromosomal recruitment of p-MDC1 (Figs. 3A and 3B). Despite impaired recruitment of p-MDC1, TUNEL signals increased after ETP treatment with nocodazole. Unexpectedly, however, TUNEL signals decreased more during recovery in the presence of nocodazole than in the absence of nocodazole (Figs. 3C and 3D). Because metaphase bivalents were disorganized and lost their conventional cruciform shape after recovery with nocodazole, we thought that this is not likely associated with DSB repair (Fig. 3C). Indeed, comet assay revealed that a substantial amount of DNA damage remained after recovery with nocodazole (Figs. S3A and 3B), suggesting that spindle microtubules are required to maintain chromosome structures during recovery from DNA damage. Consistent with this, an abnormal association of cohesin and condensin components SMC3 and SMC4 was observed in metaphase bivalents after nocodazole treatment during recovery (Figs. 3E–3H). In contrast, nocodazole treatment in intact oocytes did not disrupt the compact structure of the bivalent chromosomes (Figs. S3C–3F). Therefore, our results suggest that spindle microtubules are critical to ensure DNA repair and for maintaining the chromosome architecture (Fig. 3I).

**Fig. 3.**
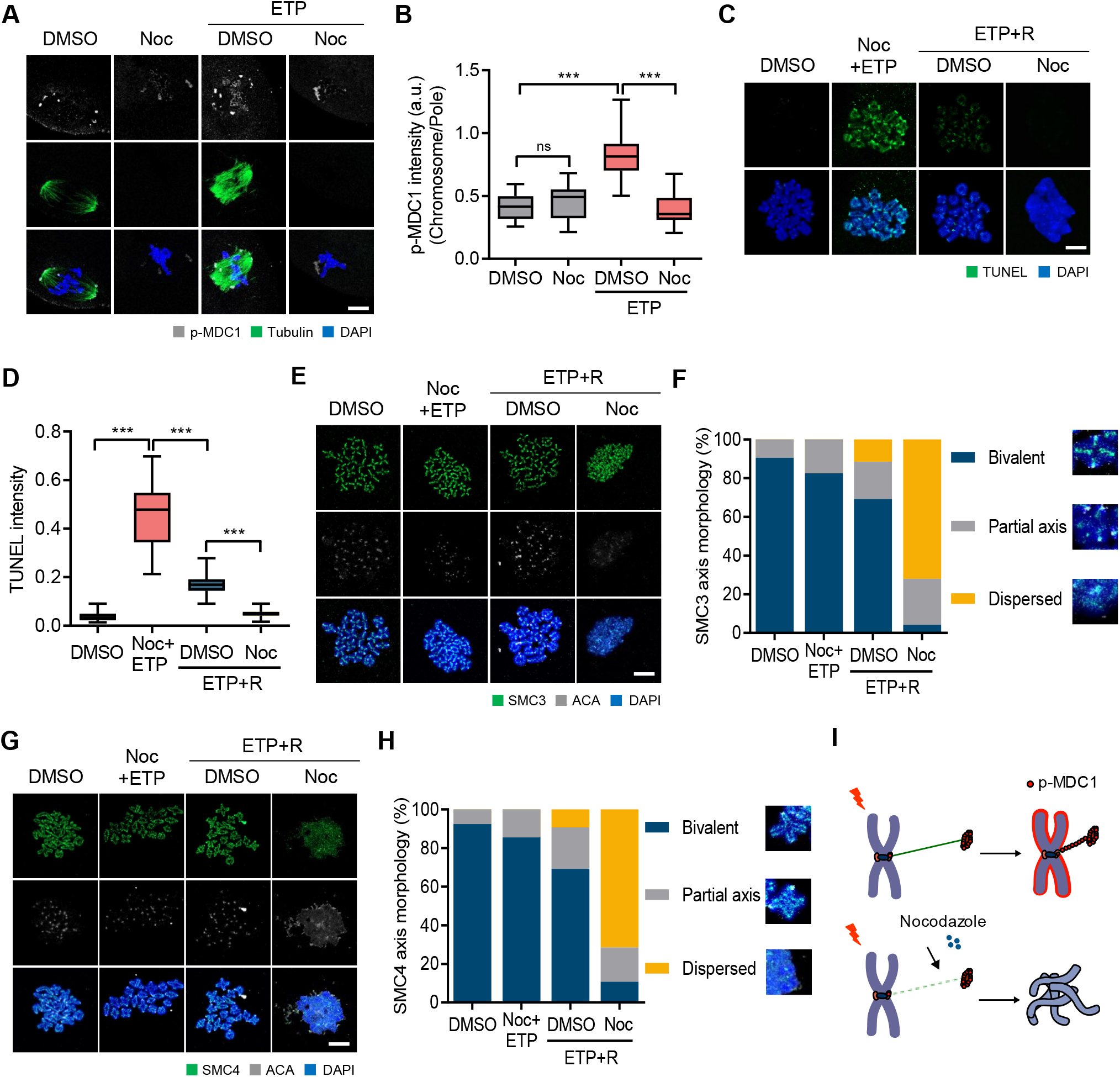
Roles of spindle microtubules in DSB repair in oocytes. (A) Representative images showing control and ETP-treated MI oocytes stained with p-MDC1 and α-tubulin antibodies after nocodazole (Noc) treatment. (B) Ratio of p-MDC1 intensity at chromosomes over spindle poles. Data are presented as the mean ± SEM from three independent experiments. ***p < 0.0001. ns, not significant. (C) Representative images of chromosome spreads showing TUNEL signals from control, ETP-treated with Noc (ETP+Noc) and recovered oocytes (ETP+R) with or without Noc. (D) Quantification of TUNEL intensity. Data are presented as the mean ± SEM from three independent experiments. ***p < 0.0001. (E, G) Representative images of chromosome spreads stained with SMC3 or SMC4 antibodies. (F, H) Quantification of SMC3 or SMC4 axis morphology. (I) Schematic illustration showing the roles of spindle microtubules in DSB repair.

### CIP2A is essential for relocating p-MDC1 and p-TOPB1 from the spindle pole to chromosomes

It was demonstrated previously that CIP2A is specifically localized at the spindle poles with weak signals on chromosomes in MI oocytes [27]. Moreover, CIP2A has recently been shown to interact with the TOPBP1 complex during mitosis in breast cancer cells [28, 29]. Given that MDC1 interacts with TOPBP1 to tether DSBs during mitosis [15], we attempted to determine whether CIP2A is associated with the chromosomal recruitment of p-MDC1 in response to DNA damage during oocyte meiosis. Consistent with previous results, CIP2A was mainly localized at the spindle poles in MI oocytes. However, after ETP treatment, CIP2A was relocated from the spindle poles to the chromosomes, similarly to p-MDC1 (Figs. 4A and 4B).

**Fig. 4.**
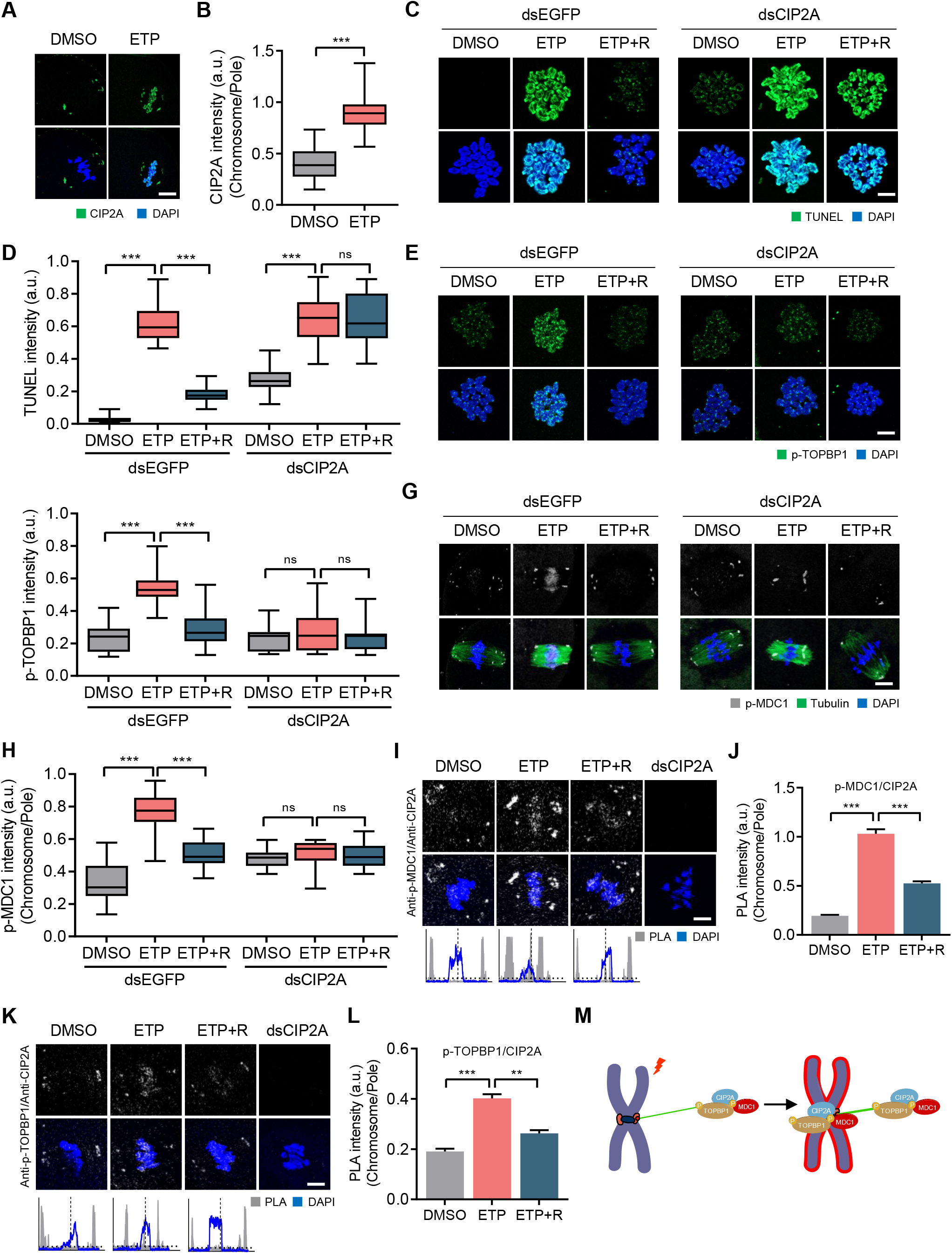
CIP2A-dependent chromosomal recruitment of p-MDC1 and p-TOPBP1. (A) Representative images of control and ETP-treated MI oocytes stained with CIP2A antibody. Scale bar, 10 μm. (B) Ratio of CIP2A intensity at chromosomes over spindle poles. Data are presented as the mean ± SEM from three independent experiments. ***p < 0.0001. (C, E, G) Representative images of chromosome spreads from control, ETP-treated, and recovered (ETP+R) oocytes, which were microinjected with either dsEGFP or dsCIP2A. TUNEL, p-TOPBP1 and p-MDC1 signals are shown. Scale bar, 10 μm. (D, F, H) Quantification of TUNEL and p-TOPBP1 intensity and ratio intensity of p-MDC1 at chromosomes over spindle poles (chromosome/pole). Data are presented as the mean ± SEM from three independent experiments. ***p < 0.0001. ns, not significant. (I, K) Representative images of PLA and profile of PLA signals between opposite spindle poles. Oocytes injected dsCIP2A were used as control. Scale bar, 10 μm. (J, L) Quantification of PLA intensity. Data are presented as the mean ± SEM from three independent experiments. **p < 0.001, ***p < 0.0001. (M) Schematic illustration showing the chromosomal recruitment of CIP2A-MDC1-TOPBP1 complex from spindle pole in response to DNA damage.

The above observations imply that CIP2A may play a role in the DDR during oocyte meiosis. To investigate this, we depleted CIP2A and assessed the DNA damage levels after ETP treatment. TUNEL signals significantly increased after CIP2A depletion, and this increase was more pronounced after ETP treatment. Notably, the TUNEL signals remained high after recovery in CIP2A-depleted oocytes, indicating that CIP2A depletion impairs DSB repair (Figs. 4C and 4D).

We next investigated the localization of p-MDC1 and p-TOPBP1 after ETP treatment in CIP2A-depleted oocytes. Consistent with previous reports that CIP2A interacts with TOPBP1 at sites of DSB in mitotic chromosomes [28, 29], p-TOPBP1 levels also increased in chromosomes after ETP treatment but decreased after recovery, similar to the chromosomal recruitment of CIP2A in response to DNA damage. However, in CIP2A-depleted oocytes, p-TOPBP1 levels did not increase in the chromosomes after ETP treatment and remained low after recovery (Figs. 4E and 4F). Similarly, the chromosomal recruitment of p-MDC1 after ETP treatment was abolished after CIP2A depletion (Figs. 4G and 4H), suggesting that CIP2A is required to recruit p-MDC1 and p-TOPBP1 on chromosomes in response to DNA damage.

To further clarify the relationship between CIP2A, p-MDC1, and p-TOPBP1, we performed an *in situ* proximity ligation assay (PLA), which can visualize the *in vivo* interaction between the two proteins [30]. Our results showed that PLA signals between CIP2A and p-MDC1 or p-TOPTBP1 mainly occurred at the spindle poles but were enriched at the chromosomes after ETP treatment. However, PLA signals at the chromosomes decreased after recovery, indicating that CIP2A interacted with p-MDC1 and p-TOPBP1 at the spindle poles and relocated to the chromosomes in response to DNA damage (Figs. 4I–4L). Therefore, along with the observation that microtubule depolymerization impaired chromosomal recruitment of p-MDC1 after ETP treatment, our results suggest that microtubule-dependent transport of the CIP2A-MDC1-TOPBP1 complex from spindle poles to chromosomes is necessary to ensure DSB repair during oocyte meiosis (Fig. 4M).

### Kinetochores and centromeres are structural hubs for chromosomal recruitment of CIP2A-MDC1-TOPBP1 complex for DSB repair

Kinetochores and centromeres serve as structural platforms that mediate the interactions between chromosomes and spindle microtubules. Therefore, we hypothesized that kinetochores and centromeres act as structural hubs for the chromosomal recruitment of the CIP2A-MDC1-TOPBP1 complex. To investigate this, we depleted CENP-A using the Trim-away method, a new technique that acutely and rapidly degrades endogenous proteins in oocytes [31, 32]. We found that CENP-A levels significantly decreased within 4 h of injecting CENP-A antibody into Trim21-expressing oocytes (Figs. S4A and S4B). Although CENP-A depletion did not change the chromosome structure, including the SMC3/4 distribution, and kMT attachments (Figs. S4C–S4H), ACA and p-MDC1 levels decreased significantly at the centromeres after CENP-A depletion (Figs. 5A and 5B). Moreover, CENP-A depletion impaired chromosomal recruitment of CIP2A and p-MDC1 after ETP treatment (Figs. 5C–5E). Consistent with the impaired recruitment of CIP2A and p-MDC1 in response to DNA damage, TUNEL signals remained high after CENP-A depletion and did not decrease after recovery (Figs. 5F and 5G). Intriguingly, the increase in TUNEL signals was more pronounced in centromeres after CENP-A depletion (Figs. 5F and 5H).

**Fig. 5.**
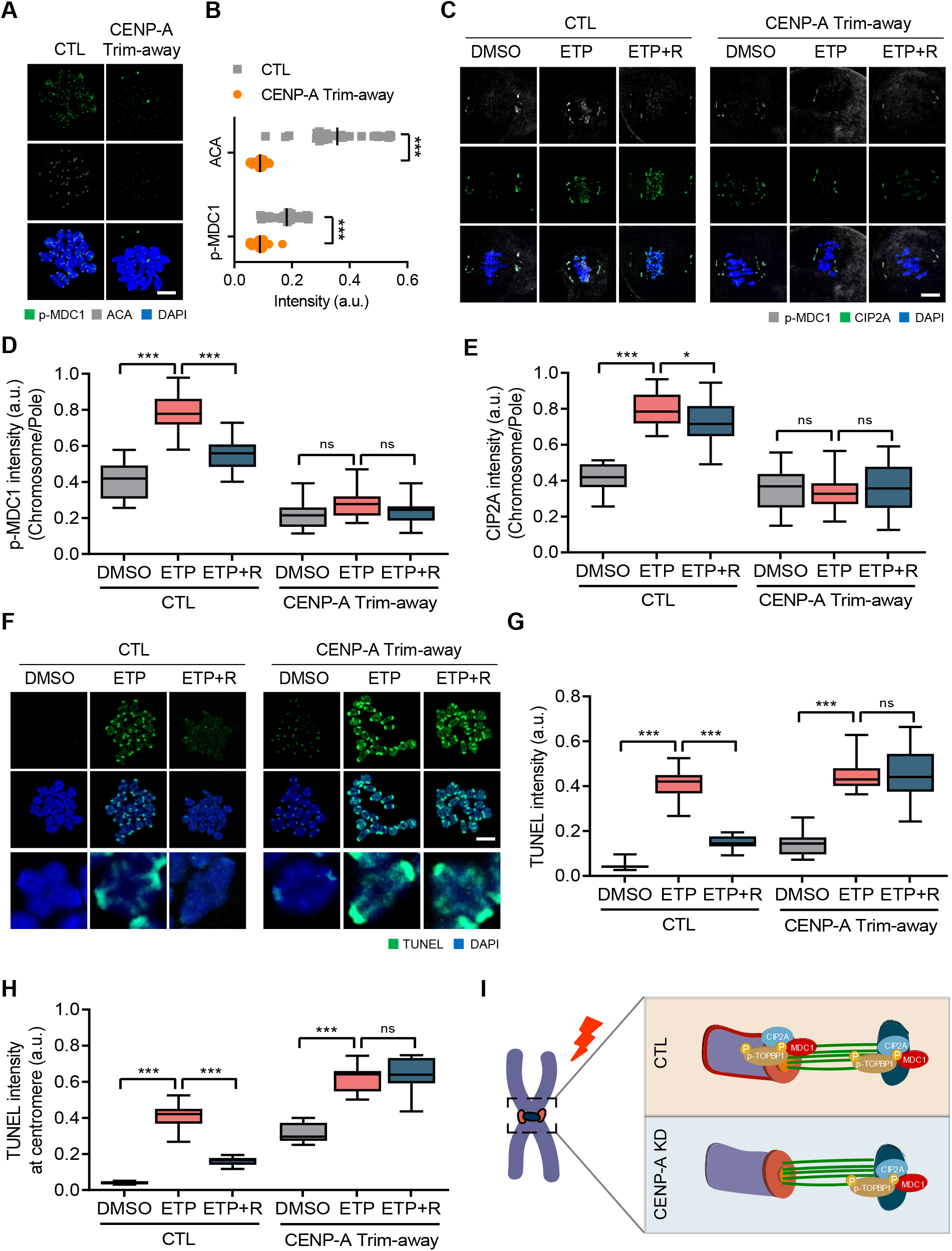
Impaired DSB repair after CENP-A depletion. (A) Representative images of control and CENP-A Trim-away oocytes stained with p-MDC1 and ACA antibodies. Scale bar, 10 μm. (B) Quantification of ACA and p-MDC1 intensities. Data are presented as the mean ± SEM from three independent experiments. ***p < 0.0001. (C) Representative images of control and CENP-A Trim-away oocytes stained with p-MDC1 and CIP2A antibodies after ETP treatment and recovery (ETP+R). Scale bar, 10 μm. (D, E) Ratio of p-MDC1 and CIP2A intensities at chromosomes over spindle poles. Data are presented as the mean ± SEM from three independent experiments. ***p < 0.0001. ns, not significant. (F) Representative images of chromosome spreads showing TUNEL signals in control and CENP-A Trim-away oocytes after ETP treatment and recovery. Scale bar, 10 μm. Representative individual chromosomes are enlarged (bottom). (G, H) Quantification of TUNEL intensity on entire chromosomes and at centromeres. Data are presented as the mean ± SEM from three independent experiments. ***p < 0.0001. ns, not significant. (I) Schematic diagram depicting the roles of CENP-A in CIP2A-MDC1-TOPBP1 chromosomal recruitment.

Similar to CENP-A depletion, kinetochore protein HEC1 depletion also impaired the recruitment of p-MDC1 and DSB repair without affecting the chromosome structure (Fig. S5). Altogether, our results demonstrate that kinetochores and centromeres serve as a structural hub for microtubule-dependent chromosomal recruitment of the CIP2A-MDC1-TOPBP1 complex for DSB repair during oocyte meiosis (Fig. 5I).

### Chromosomal relocation of CIP2A-MDC1-TOPBP1 is regulated by PLK1 activity

It is well established that ATM is a key kinase that regulates DDR [2]. Moreover, emerging evidence suggests that PLK1 is inactivated in response to DNA damage in mitotic cells [33–35]. Therefore, we investigated the roles of ATM and PLK1 in the chromosomal relocation of the CIP2A-MDC1-TOPBP1 complex during oocyte meiosis. After treating MI oocytes with either BI2536 (PLK1 inhibitor) or KU55933 (ATM inhibitor), we examined changes in CIP2A and p-MDC1 localization after ETP treatment. Our data showed that CIP2A and p-MDC1 signals disappeared at the spindle poles and were enriched on chromosomes after BI2536 treatment. These chromosomal signals increased further after ETP treatment and did not return to the spindle poles after recovery (Figs. S6A–6C). In contrast to PLK1, ATM inhibition did not impair the chromosomal relocation of CIP2A and p-MDC1 after ETP treatment (Figs. S6D–6F), suggesting that DNA damage-induced CIP2A-MDC1-TOPBP1 relocation is regulated by PLK1 activity but not by ATM activity.

Because PLK1 inhibition led to CIP2A relocation to chromosomes, we hypothesized that DSBs could induce PLK1 inactivation at the spindle poles, which in turn causes chromosomal recruitment of CIP2A in association with MDC1 and TOPBP1. To test this hypothesis, we examined the phosphorylated PLK1 at T210 (p-T210-PLK1) after ETP treatment. We found that p-PLK1 signals were mainly detectable at spindle poles and chromosomes in intact oocytes. After ETP treatment, however, p-PLK1 signals significantly decreased at spindle poles but not on chromosomes, which were rescued after 2 h of recovery (Figs. 6A and 6B). To further investigate the role of PLK1 in the chromosomal relocation of the CIP2A-MDC1-TOPBP1 complex and subsequent DSB repair during oocyte meiosis, we overexpressed a constitutively active PLK1-T210D mutant in oocytes. The overexpression of the PLK1-T210D mutant disrupted the pole-specific decrease in p-PLK1 levels after ETP treatment (Figs. S6G and S6H). The PLK1-T210D overexpression also impaired the chromosomal relocation of CIP2A and p-MDC1 after ETP treatment (Figs. 6C–6E). Consistent with this finding, TUNEL signals did not decrease after recovery in oocytes overexpressing PLK1-T210D (Figs. 6F and 6G). Therefore, our results suggest that PLK1 inactivation in response to DNA damage at the spindle poles is a prerequisite for chromosomal relocation of the CIP2A-MDC1-TOPBP1 complex to ensure DSB repair during oocyte meiosis.

**Fig. 6.**
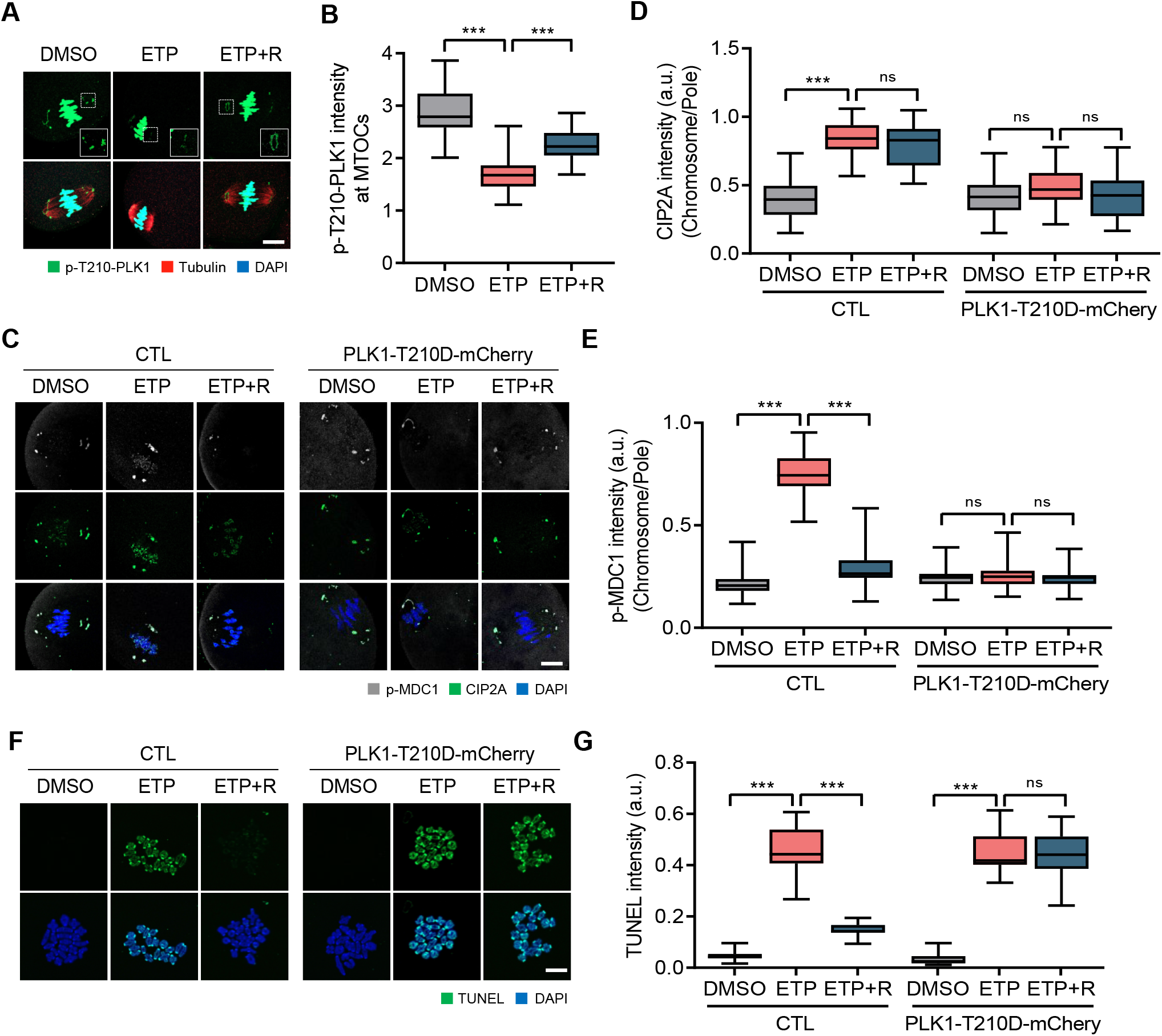
DSB-induced PLK1 inactivation at spindle poles. (A) Representative images of oocytes showing localization of p-T210-PLK1 after ETP treatment and recovery (ETP+R). Scale bar, 10 μm. (B) Quantification of p-T210-PLK1 intensity at MTOCs. Data are presented as the mean ± SEM from three independent experiments. ***p < 0.0001. (C) Representative images showing p-MDC1 and CIP2A signals in oocytes expressing either mCherry or PLK1-T210D-mCherry. Scale bar, 10 μm. (D, E) Ratio of CIP2A and p-MDC1 intensities at chromosomes over spindle poles. Data are presented as the mean ± SEM from three independent experiments. ***p < 0.0001. ns, not significant. (F) Representative images of chromosome spreads showing TUNEL signals in oocytes expressing either mCherry or PLK1-T210D-mCherry. Scale bar, 10 μm. (G) Quantification of TUNEL intensity. Data are presented as the mean ± SEM from three independent experiments. ***p < 0.0001. ns, not significant.

### Dephosphorylation of CIP2A at S904 triggers chromosomal relocation of the CIP2A-MDC1-TOPBP1 complex from the spindle pole

It was shown previously that CIP2A is phosphorylated by PLK1 at S904, which targets CIP2A in MTOCs and facilitates MTOC organization in association with CEP192 during oocyte meiosis [27]. Because PLK1 is inactivated at the spindle poles after DNA damage, we sought to investigate whether DNA damage-induced PLK1 inactivation at the spindle pole causes dephosphorylation of CIP2A at S904, which in turn induces CIP2A relocation to chromosomes. To test this possibility, we depleted endogenous CIP2A and ectopically expressed the CIP2A-S904A and CIP2A-S904D mutants. While the CIP2A-S904D mutant exhibited MTOC (spindle pole) localization similar to that of endogenous CIP2A, the CIP2A-S904A mutant localized mainly on the chromosomes (Figs. 7A–7F). The MTOC and chromosomal localization of CIP2A-S904D and CIP2A-S904A, respectively, did not change after ETP treatment (Figs. 7A–7F; Fig. S7). Similar to CIP2A localization, p-MDC1 signals remained on the chromosomes after ETP treatment and did not return to the spindle poles after recovery in oocytes expressing CIP2A-S904A (Figs. 7A and 7B). In contrast, in oocytes expressing CIP2A-S904D, p-MDC1 signals did not appear on the chromosomes after ETP treatment (Figs. 7D and 7E), implying that the S904 residue at CIP2A is a *bona fide* phosphorylation site that regulates CIP2A relocation. Moreover, TUNEL signals revealed that oocytes expressing CIP2A-904D did not repair DSBs after recovery, whereas oocytes expressing CIP2A-S904A partially repaired DSBs (Figs. 7G and 7H), indicating that chromosomal recruitment of CIP2A is essential for DSB repair. Collectively, our results suggest that DSB-induced PLK1 inactivation at the spindle poles causes CIP2A dephosphorylation at S904, which in turn promotes chromosomal relocation of CIP2A along with p-MDC1 and p-TOPBP1 to ensure DSB repair.

**Fig. 7.**
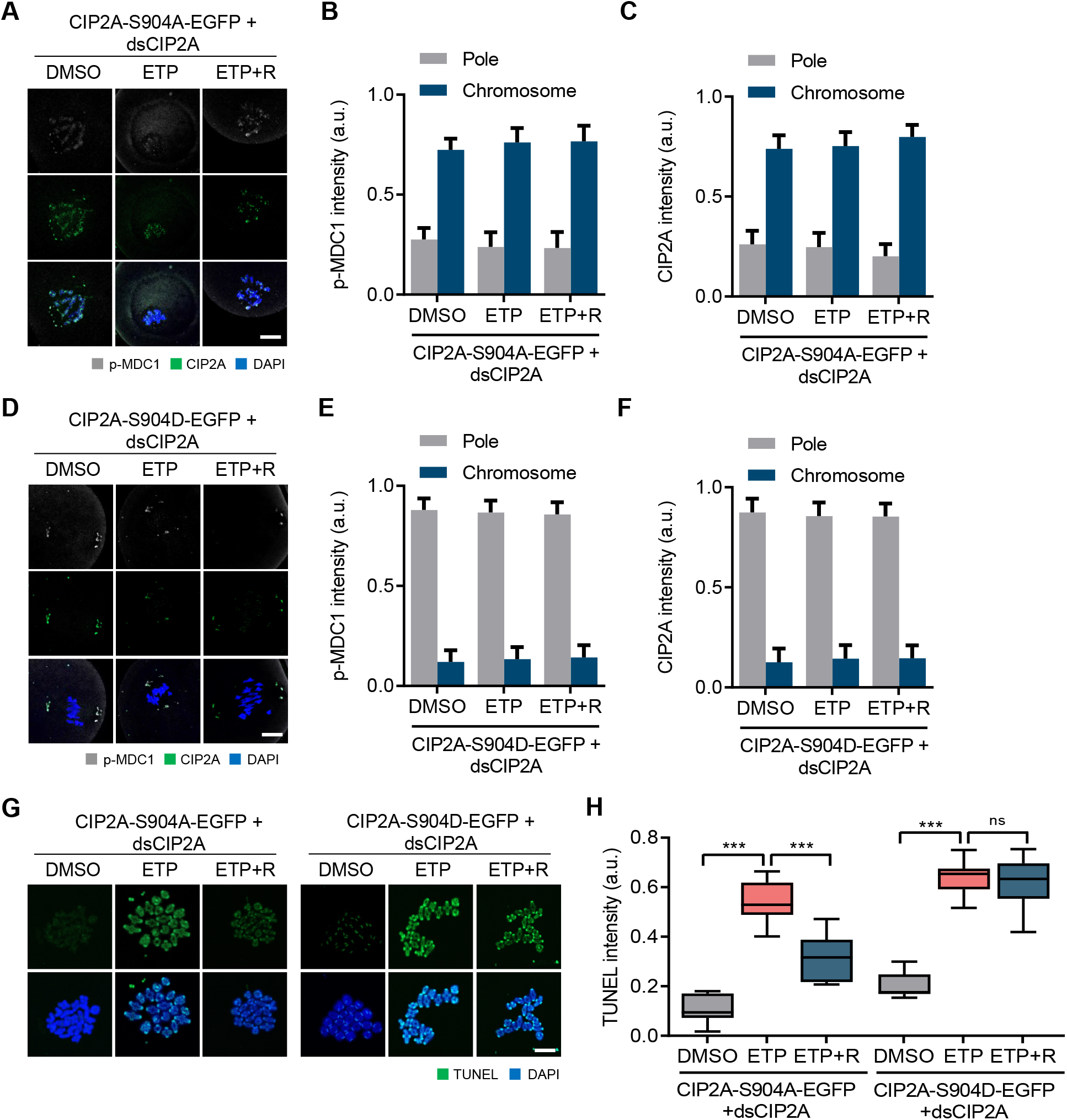
CIP2A dephosphorylation by PLK1 inactivation after DNA damage. (A, D) Representative images showing CIP2A and p-MDC1 signals in oocytes co-injected with dsCIP2A and mRNAs encoding either CIP2A-S904A-EGFP or CIP2A-S904D-EGFP. Scale bar, 10 μm. (B, C, E, F) Quantification of CIP2A and p-MDC1 localization (chromosomes vs pole). Data are presented as the mean ± SEM from two independent experiments. ***p < 0.0001. ns, not significant. (G) Representative images showing TUNEL signals in oocytes co-injected with dsCIP2A and mRNAs encoding either CIP2A-S904A-EGFP or CIP2A-S904D-EGFP. Scale bar, 10 μm. (H) Quantification of TUNEL intensity. Data are presented as the mean ± SEM from two independent experiments. ***p < 0.0001. ns, not significant.

## Discussion

In this study, we uncovered a previously unknown pathway of DDR that occur during meiosis in mouse oocytes. After DNA damage, the spindle microtubules rapidly shrink and are stabilized. Moreover, DSBs induce PLK1 inactivation at the spindle poles, in turn causing CIP2A dephosphorylation at S904. Once dephosphorylated, CIP2A complexed with p-MDC1 and p-TOPBP1 at the spindle poles moves along spindle microtubules and is recruited to chromosomes through kinetochores and centromeres. This chromosomal relocation of the CIP2A-MDC1-TOPBP1 complex further ensures the recruitment of downstream repair factors, including BRCA1 and 53BP1, and the subsequent DNA repair during oocyte meiosis (Fig. 8).

**Fig. 8.**
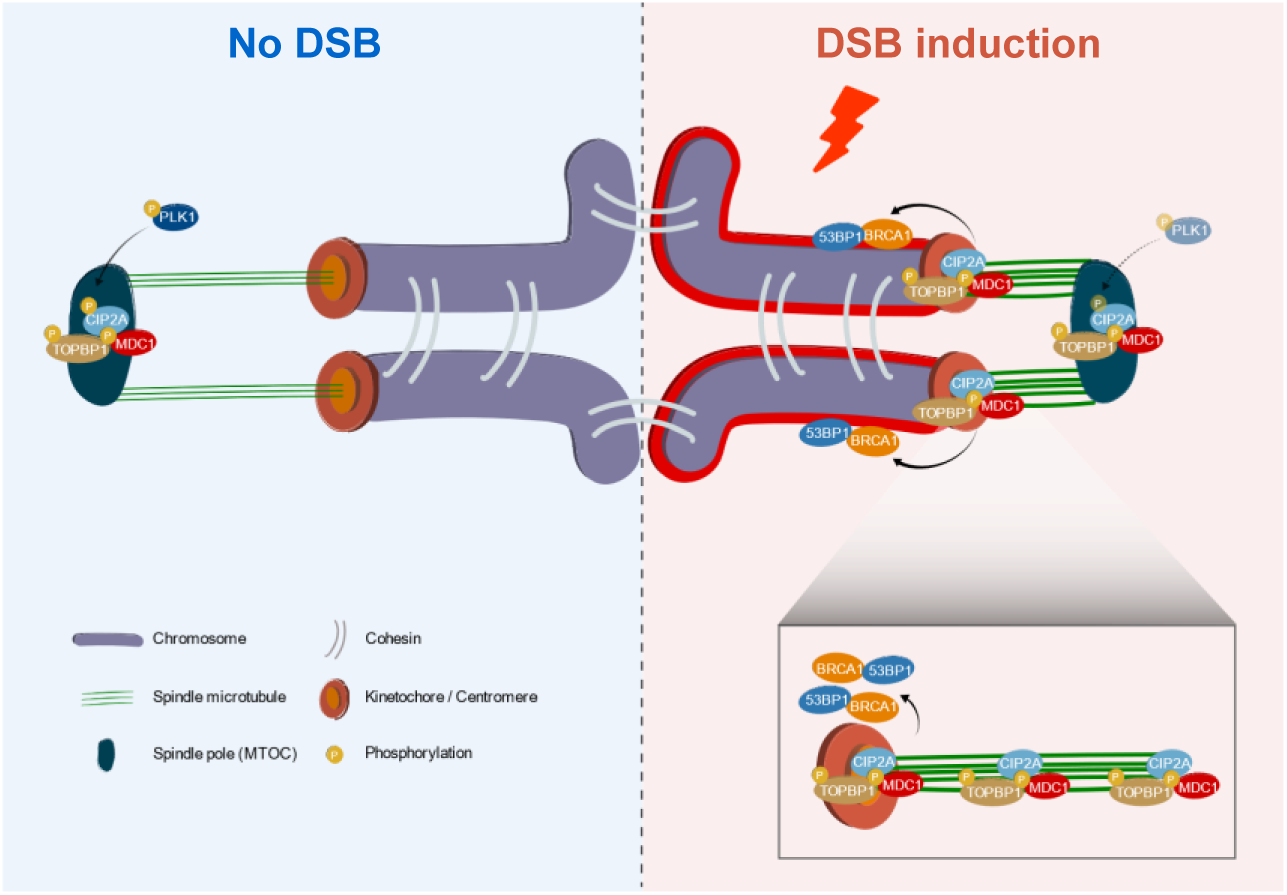
Schematic model of DNA damage response that occurs in oocytes during meiosis I. After DNA damage, spindle microtubules rapidly shrink and are stabilized. Moreover, DSBs induce the PLK1 inactivation at spindle poles, which in turn causes CIP2A dephosphorylation. Once dephosphorylated, CIP2A complexed with p-MDC1 and p-TOPBP1 at spindle poles moves along spindle microtubules and is recruited to chromosomes through kinetochores and centromeres, which further ensures the recruitment of downstream repair factors, including BRCA1 and 53BP1, and subsequent DNA repair during oocyte meiosis.

Mitotic cells can efficiently mark DSB sites by tethering broken ends with MDC1 and TOPBP1, but downstream repair factors such as RNF8, BRCA1 and 53BP1, are not recruited to DSB sites during mitosis [11, 14, 15]. This is associated with mitosis-specific phosphorylation of RNF8 and 53BP1. CDK1 and PLK1 phosphorylate 53BP1 on T1609 and T1618, which prevents its binding to γ-H2AX [36, 37]. CDK1 also phosphorylates RNF8 on T198 and prevents its interaction with MDC1 and recruitment to DSB sites [16]. In contrast to mitotic cells, our data show that oocytes can recruit BRCA1 and 53BP1 and repair DSBs during meiosis. Although the mechanisms that allow the recruitment of BRCA1 and 53BP1 to DSBs in oocytes remain elusive, given that oocytes are enormous and spend a long time in meiosis, one can speculate that DNA damage temporally and/or spatially inactivates CDK1 or PLK1 in oocytes, unlike in mitotic cells. Indeed, it was observed that PLK1 was inactivated at the spindle pole of oocytes after DNA damage. Consistent with this, emerging evidence suggests that PLK1 is inactivated in response to DNA damage in mitotic cells [33–35]. Therefore, it is possible that mitosis-specific phosphorylations of RNF8 and 53BP1 could be temporarily dampened after DNA damage in oocytes during meiosis. Despite the critical difference from mitosis, meiosis shares mechanisms and regulation with mitosis in many respects. Therefore, we cannot entirely exclude the possibility that the DSB repair observed during oocyte meiosis may occur in mitotic cells under certain conditions. Consistent with this notion, evidence supporting that mitotic cells are capable of DNA repair is only beginning to emerge [38–40]. Given that oocytes and cancer cells have certain similarities, including a certain degree of aneuploidy, the ability to sort and cluster MTOCs, and cortical softening for cell division [41], it is also noteworthy that mitotic DDR is frequently observed in many cancer cells [13]. Further studies are necessary to clarify the occurrence of DSB repair during mitosis.

Spindle microtubules exhibit highly dynamic instability, repeating the cycle of growth and shrinkage. This fast turnover of spindle microtubules is thought to help correct the erroneous attachments that occur during bipolar chromosome alignment. In this study, we demonstrated that spindle microtubules are shortened and stabilized in response to DNA damage during oocyte meiosis. The biological significance of spindle shrinkage and stabilization remains unclear, but it is notable that many proteins involved in DDR pathways are associated with spindle microtubules and mitotic proteins [42]. Considering that not all DDR factors found in the nucleus are bound to chromosomes in the absence of DNA damage, spindle association of these factors after nuclear envelope breakdown is not surprising. In this regard, one can speculate that spindle shrinkage and stabilization induced by DSBs are a strategy to expedite DNA repair by placing DDR factors in close proximity to facilitate chromosomal recruitment. Furthermore, spindle shrinkage and stabilization still occurred either after inhibiting ATM or PLK1 activity or after depleting CENP-A or HEC1. This suggests that ATM, PLK1, and kMT attachments are not the driving forces regulating spindle changes in response to DNA damage. Given the close communication between actin filaments and microtubules in spindle assembly in oocytes [43], it is reasonable to postulate that actin filaments are involved in the process governing DSB-induced spindle changes. Indeed, DNA damage has been shown to induce actin filament assembly to promote efficient DNA repair [44].

We found that MDC1 and TOPBP1 levels were significantly reduced after CIP2A depletion. Moreover, the PLA signals showed that CIP2A formed a triple complex with MDC1 and TOPBP1 at the spindle poles. Thus, it is likely that the CIP2A-MDC1-TOPBP1 complex plays an important role in maintaining MTOCs at the spindle pole, and, when DSBs are induced, PLK1 is selectively inactivated at the spindle pole, which in turn causes CIP2A dephosphorylation. Because comparable amounts of PLA signals were detectable at the spindle pole and chromosomes after DNA damage, CIP2A dephosphorylation does not seem to affect complex formation with MDC1 and TOPBP1 and is thought to be involved only in the transport of CIP2A. Therefore, our results suggest that chromosomal relocation of MDC1-TOPBP1 for DNA repair is CIP2A-dependent and that dephosphorylation of CIP2A acts as a trigger for chromosomal transport of the CIP2A-MDC1-TOPBP1 complex in response to DNA damage.

Our data revealed that kinetochores and centromeres act as structural hubs for the microtubule-dependent pole-to-chromosome relocation of the CIP2A-MDC1-TOPBP1 complex. Because CENP-A is remarkably stable and displays spectacular longevity in mouse oocytes [45], we depleted CENP-A using the Trim-away method. Consistent with a previous report that CENP-A is dispensable for mitotic centromere function after the initial centromere/kinetochore assembly [46], CENP-A depletion did not affect the overall architecture of bivalent chromosomes and kMT attachments in mouse oocytes. However, chromosomal recruitment of CIP2A and p-MDC1 after DNA damage was severely impaired after CENP-A depletion, despite comparable levels of CIP2A and p-MDC1 at the spindle poles. To further clarify the role of centromeres in chromosomal recruitment of the CIP2A-MDC1-TOPBP1 complex, we depleted HEC1 and obtained similar results seen after CENP-A depletion. Therefore, it is proposed that the kinetochore and centromere act as a structural hub to recruit the CIP2A-MDC1-TOPBP1 complex from the spindle poles, implying crosstalk between the centromere/kinetochore and spindle poles in response to DNA damage. This mutual crosstalk is supported by a recent finding that centromere dysfunction compromises mitotic spindle pole integrity [47]. Considering that centromeres are fragile and sensitive to DNA damage because of their repetitive sequences, one can speculate that centromeres rapidly recognize DNA damage and then deliver damage signals to the spindle poles through microtubules. However, this crosstalk between the centromere/kinetochore and spindle poles seems to be different from that inducing spindle changes after DNA damage because DSB-induced spindle shrinkage still occurred after depleting CENP-A or HEC1. Our findings provide a strong foundation for future mechanistic studies to elucidate how chromosomes crosstalk with spindle poles in response to DNA damage.

In summary, our data reveal that oocytes can repair DSBs during meiosis I through the microtubule-dependent chromosomal recruitment of CIP2A-MDC1-TOPBP1 from the spindle pole via the centromere and kinetochores. These data provide new insights into the critical crosstalk between chromosomes and spindle microtubules in response to DNA damage to maintain genomic stability during oocyte meiosis.

## Materials and methods

All chemicals and reagents were obtained from Sigma-Aldrich unless otherwise stated.

### Oocytes collection and culture

All procedures for mouse care and use were conducted in accordance with the guidelines of and were approved by the Institutional Animal Care and Use Committee of Sungkyunkwan University (approval ID: SKKUIACUC-2022-06-24-2). Female CD1 mice were purchased from a local company (Koatech, Korea). To collect oocytes, three-to four-week-old female mice were intraperitoneally injected with 5 IU of pregnant mare serum gonadotropin (PMSG). After 46–48 h, fully grown GV oocytes were collected from the follicles and recovered in M2 medium (M7167) supplemented with 100 μM of 3-isobuthly-1-methlxanthine (IBMX; I5897). To obtain oocytes at the MI stage, GV oocytes were cultured *in vitro* in IBMX-free M2 medium for 8 h in mineral oil (M5310) at 37°C in a 5% CO_2_ atmosphere.

### Chemical treatment

To induce DSBs, MI oocytes were exposed to 50 μg/mL etoposide (ETP) for 30 min. For recovery, the ETP-treated oocytes were washed and cultured in a fresh M2 medium for 2 h. To depolymerize spindle microtubules, MI oocytes were treated with 20 μg/mL nocodazole (SML1665) for 10 min. For ATM and PLK1 inhibition, oocytes were treated with 10 μM ATM inhibitor (KU55933, Selleckchem) and 200 nM PLK1 inhibitor (BI2536, Selleckchem), respectively. Control oocytes were treated with dimethyl sulfoxide (DMSO).

For the analysis of kMT attachment, MI oocytes were placed in an ice-cold M2 medium for 10 min prior to immunostaining.

### Overexpression of CIP2A and PLK1 mutants

CIP2A-S904A and CIP2A-S904D were subcloned into pRN3-mCherry vectors, as described previously [27]. PLK1-T210D clones were obtained from Addgene (#68133) and were subcloned into the pRN3-mCherry vector. PLK1 mRNAs were transcribed *in vitro* using a T3 mMESSAGE mMACHINE (Ambion) according to the manufacturer’s protocol. After purification, mRNAs were diluted to a concentration of approximately 500 ng/μL and microinjected into GV oocytes on the stage of a DMIRB inverted microscope (Leica) using a FemtoJet microinjector (Eppendorf) and micromanipulators (Narishige).

### Depletion of CIP2A, CENP-A, and HEC1

The CIP2A depletion was performed as described previously [27]. Briefly, GV oocytes were microinjected with double-stranded RNA targeting the endogenous CIP2A (dsCIP2A) mRNA. Control oocytes were injected with double-stranded EGFP (dsEGFP) mRNA. Microinjection was performed as previously described [48]. After microinjection, GV oocytes were maintained in an M2 medium containing IBMX for 24 h.

For CENP-A or HEC1 depletion using the Trim-away method, GV oocytes were injected with mRNA encoding Trim21-mCherry, as described previously [49]. After 1 h of culture in IBMX-containing M2 medium to allow protein expression, oocytes were cultured in an IBMX-free medium for 4 h and then microinjected with CENP-A or HEC1 antibodies at a final concentration of 100 ng/μL. Control oocytes were injected with normal IgG antibodies (sc-2025; Santa Cruz Biotechnology). Following an additional 4 h of culture after antibody injection, oocytes were subjected to immunostaining or chromosome spreading.

### Immunostaining

The oocytes were fixed with 4% paraformaldehyde in PBS for 10 min and permeabilized with 0.1% Triton X-100 and 0.01% Tween-20 for 20 min at room temperature. After blocking with 3% BSA-supplemented PBS for 1 h, the oocytes were incubated with primary antibodies overnight at 4°C. The primary antibodies used in this study were anti-BRCA1 (1:250, SAB2702136), anti-53BP1 (1:250, ab36823, Abcam), anti-acetylated α-tubulin (1:1000, T7451), anti-γ-H2AX (1:250, ab22551, Abcam), anti-MDC1 (1:250, ab241048, Abcam), anti-p-MDC1 (1:250, ab35967, Abcam), anti-p-TOPBP1 (1:250, AP3774a-EV, ABCEPTA), anti-SMC3 (1:100, ab128919, Abcam), anti-SMC4 (1:100, NBP1-86635, Novus Biologicals), anti-centromere (1:100, 15-234, Antibodies Incorporated), anti-CIP2A (1:500, sc-80662, Santa Cruz), anti-p-T210-PLK1 (1:250, ab39068, Abcam), anti-γ-tubulin (1:250, ab11316, Abcam), and anti-GFP (1:500, ab1218, Abcam). After being washed three times, the oocytes were incubated with secondary antibodies at room temperature for 2 h. For the secondary antibodies, Alexa Fluor-conjugated 488 (1:500, 115-545-146, Jackson ImmunoResearch), Alexa Fluor-conjugated 594 (1:500, 115-585-044, Jackson ImmunoResearch), and rhodamine (TRITC)-conjugated anti-human (Jackson ImmunoResearch, 109-025-088, 1:100) were used. After counterstaining with DAPI, the oocytes were mounted on glass slides and observed under an LSM 900 laser scanning confocal microscope (Zeiss) with a C-Apochromat 63× /1.2 oil immersion objective.

### Proximity ligation assay

Proximity ligation assay (PLA) was performed using the *in situ* detection reagent Orange kit Mouse/Rabbit (DUO92007). Mouse anti-CIP2A and rabbit anti-p-MDC1 or rabbit anti-p-TOPBP1 antibodies were conjugated with PLA PLUS and PLA MINUS probes, respectively. CIP2A-depleted oocytes were used as negative controls. The PLA signals were visualized using an LSM 900 laser scanning confocal microscope (Zeiss).

### Chromosome spreads

Chromosome spreads were prepared as previously described [26, 50]. Briefly, MI oocytes were exposed to acidic Tyrode’s solution (pH 2.5) for 2–3 min to remove the zona pellucida. After brief recovery in fresh medium, MI oocytes were fixed in 1% paraformaldehyde in distilled water (pH 9.2) containing 0.15% Triton X-100 and 3 mM of dithiothreitol. The slides were dried slowly in a humid chamber for several hours and then blocked with 1% BSA in PBS for 1 h at room temperature. The oocytes were incubated with a primary antibody overnight at 4°C and then with a secondary antibody for 2 h at room temperature. The DNA was stained with DAPI, and the slides were mounted for observation using LSM 900 laser scanning confocal microscopy.

### TUNEL assay

The TUNEL assay was performed using the *In Situ* Cell Death Detection Kit (Roche) according to the manufacturer’s instructions. Paraformaldehyde-fixed oocytes were washed three times with PBS and permeabilized with 0.15% Triton-X100 and 0.1% sodium citrate for 1 h on ice. The oocytes were washed and incubated with fluorescent-conjugated terminal deoxynucleotide transferase dUTP for 2 h at 37°C. After being washed three times, the oocytes were mounted on glass slides after counterstaining with DAPI, and the fluorescence signal was detected using an LSM 900 laser scanning confocal microscope (Zeiss).

### Comet assay

The comet assay was performed using an Alkaline CometAssay kit (Trevigen) according to the manufacturer’s instructions. Briefly, oocytes were mixed with melted agarose, placed on comet slides, and subjected to electrophoresis. The comet signals were visualized by staining with SYBR green (Invitrogen), and images were captured with a confocal microscope.

### Quantification of fluorescence intensity

All images were acquired at pixel dimensions of 1024 × 1024 and are shown as the maximum intensity of the Z-projections using an LSM 900 laser scanning confocal microscope (Zeiss). For the measurement of immunofluorescence intensity, images were captured with the same laser power, and the mean intensity of the fluorescence signals was measured and displayed in arbitrary units (a.u.). The data were analyzed using ZEN 3.4 Blue (Zeiss) and ImageJ software (National Institutes of Health) under the same processing parameters.

### Statistical analysis

All statistical analyses were performed using GraphPad Prism 9.0 (GraphPad Software). The data are representative of at least three independent experiments unless otherwise specified, and each experimental group included at least 15 oocytes. The differences between two groups were analyzed using Student’s t-test, and comparisons between more than two groups were analyzed using one-way analysis of variance (ANOVA) with Tukey’s *post hoc* test. The percentages of maturation were analyzed using arcsine transformation. P < 0.05 was considered statistically significant.

## Data Availability

All data needed to evaluate the conclusions of this study are presented in the article and/or the Supplementary Materials. Additional data related to this study can be requested from the authors.

## Acknowledgments

We appreciate and acknowledge Hayeon Kim for completing the pilot tests.

## Author contributions

JL and JSO conceived and designed the experiments. JL performed all the experiments. JL and JSO analyzed and interpreted the data. JSK provided critical feedback regarding the manuscript. JSO supervised the study. JL and JSO wrote the manuscript.

## Supplementary Figures

**Fig. S1.**
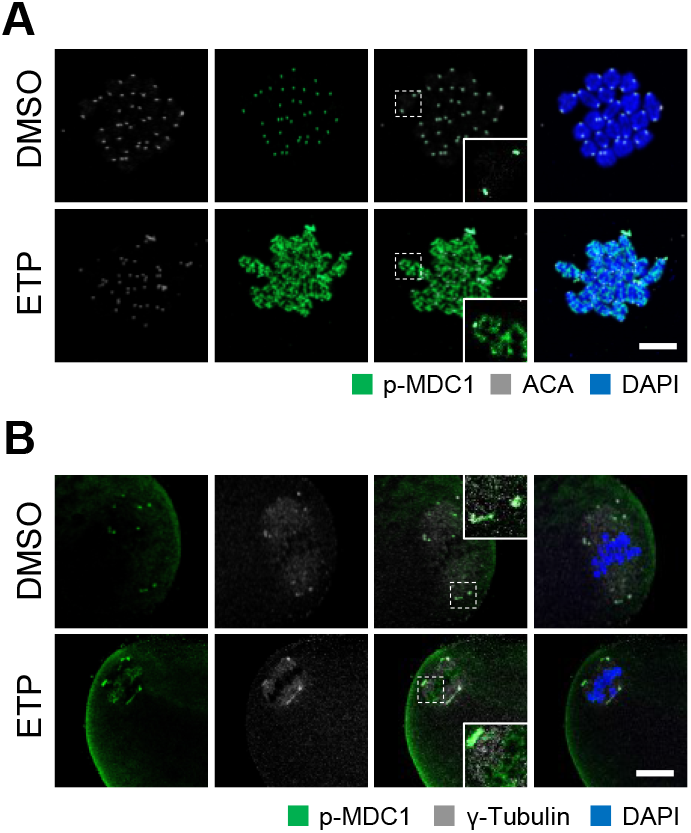
Localization of p-MDC1 after DNA damage. (A) Representative images of chromosome spreads from control and ETP-treated oocytes stained with p-MDC1 and ACA antibodies. (B) Representative images of control and ETP-treated oocytes stained with p-MDC1 and γ-tubulin antibodies. Scale bar, 10 μm.

**Fig. S2.**
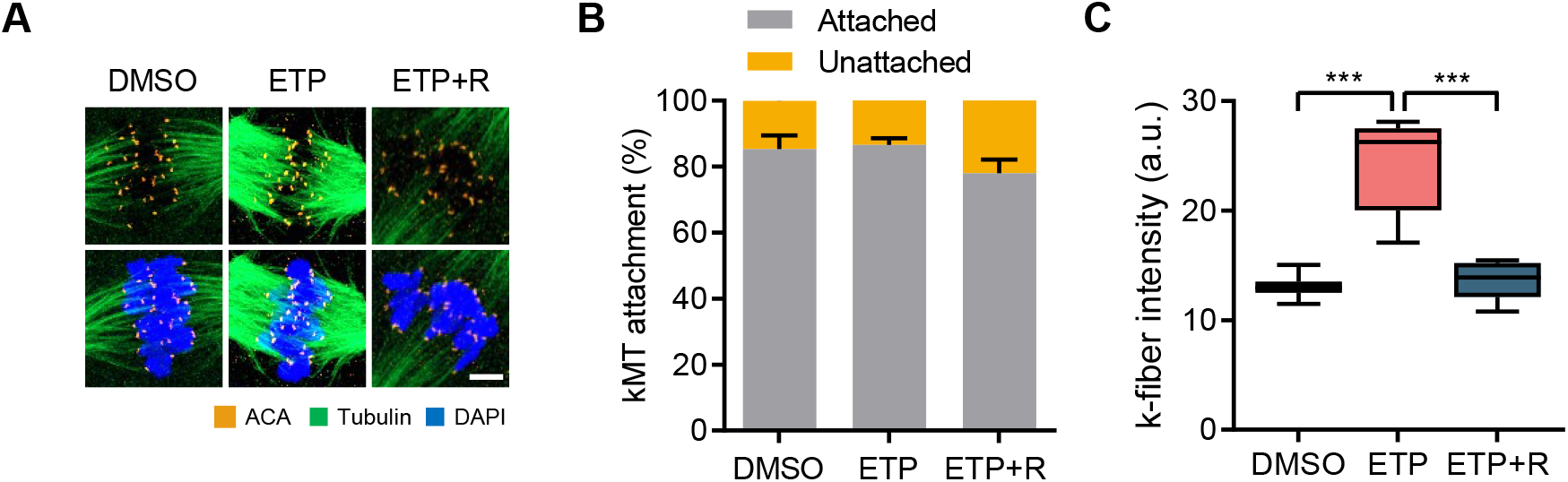
Effect of DNA damage on kMT attachment. (A) Representative images of control, ETP-treated, and recovered (ETP+R) oocytes stained with tubulin and ACA antibodies after cold treatment. Scale bar, 10 μm. (B, C) Quantification of kMT attachment and k-fiber intensity. Data are presented as the mean ± SEM from three independent experiments. ***p < 0.0001.

**Fig. S3.**
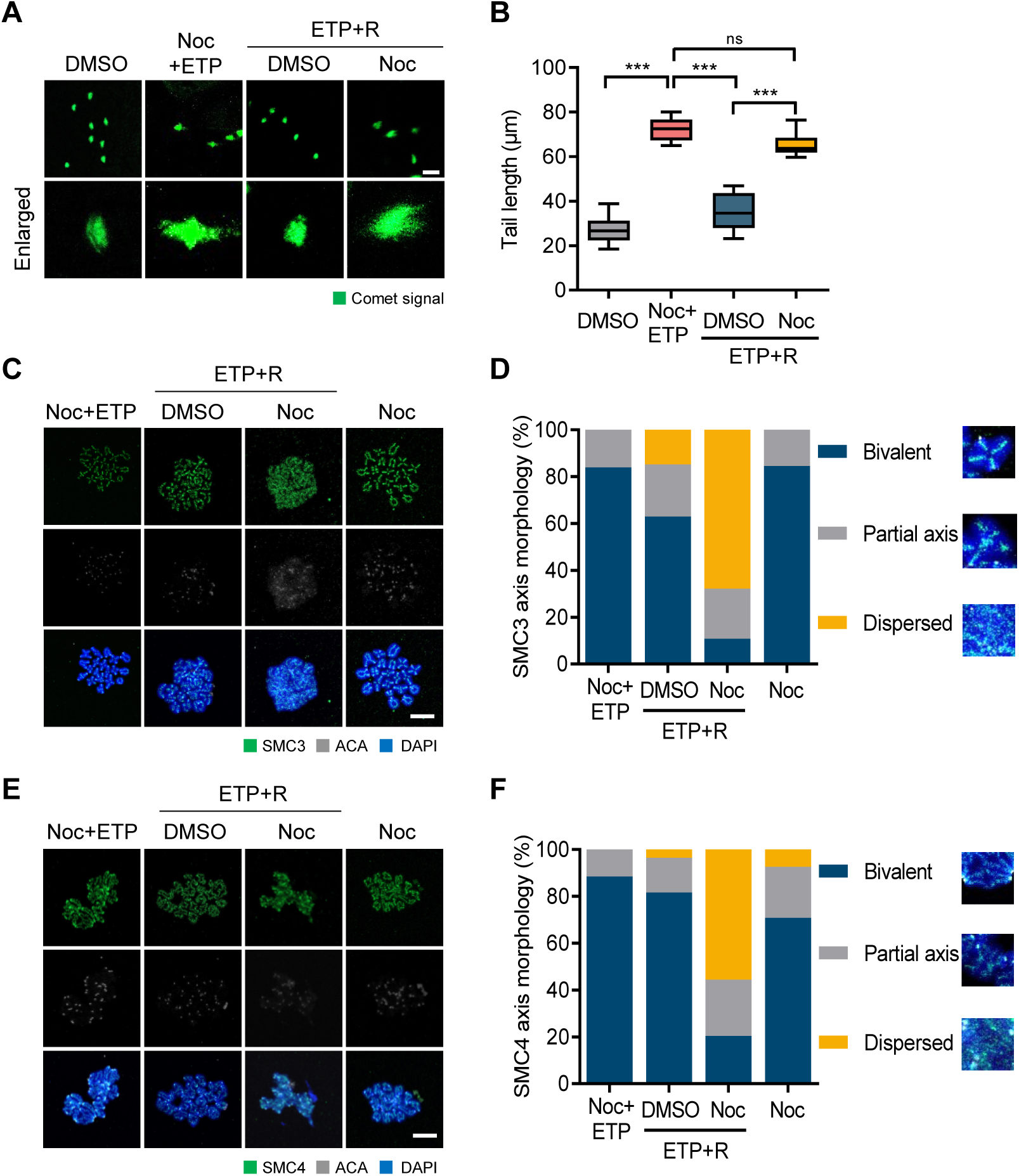
Role of spindle microtubules in maintaining chromosome integrity during DSB repair. (A) Representative images showing comet signals after nocodazole (Noc) treatment during recovery. Scale bar, 50 μm. (B) Quantification of the tail length of the comet. ***p < 0.0001, ns, not significant. (C, E) Representative images of chromosome spreads stained with SMC3 or SMC4 antibodies. Scale bar, 10 μm. MI oocytes were treated with ETP with Noc for 30 min and recovered in fresh medium for 2 h in the presence of absence of Noc. Control oocytes were treated with Noc for 2.5 h. (D, F) Quantification of SMC3 or SMC4 axis morphology.

**Fig. S4.**
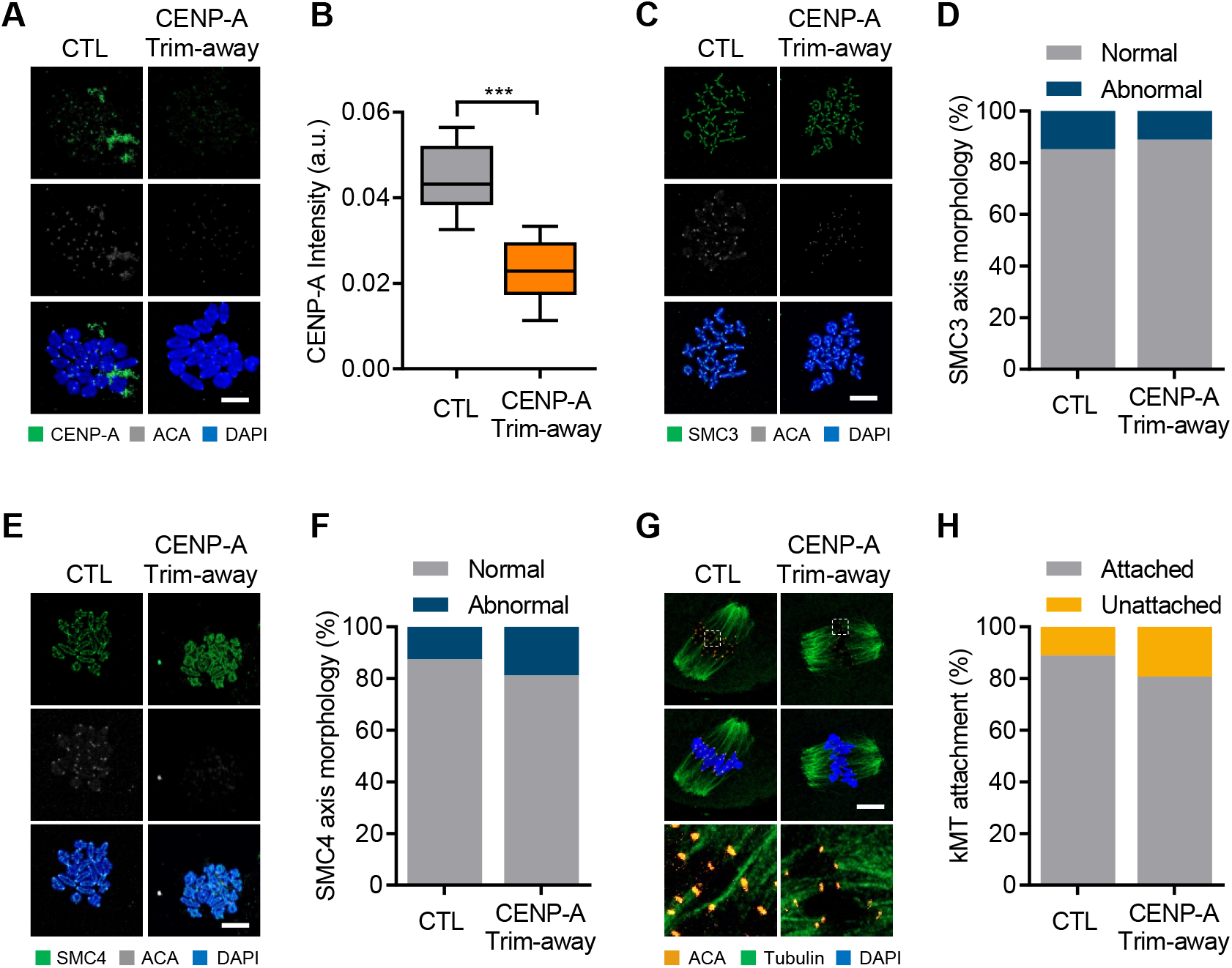
Chromosome architecture after CENP-A depletion. (A, C, E) Representative images of chromosome spreads from control and CENP-A Trim-away oocytes. Scale bar, 10 μm. (B) Quantification of CENP-A intensity. Data are presented as the mean ± SEM from three independent experiments. ***p < 0.0001. (D, F). Quantification of SMC3 orSMC4 axis morphology. (G) Representative images of control and CENP-A Trim-away oocytes stained with tubulin and ACA antibodies after cold treatment. Scale bar, 10 μm. (H) Quantification of kMT attachment. Data are presented as the mean ± SEM from three independent experiments.

**Fig. S5.**
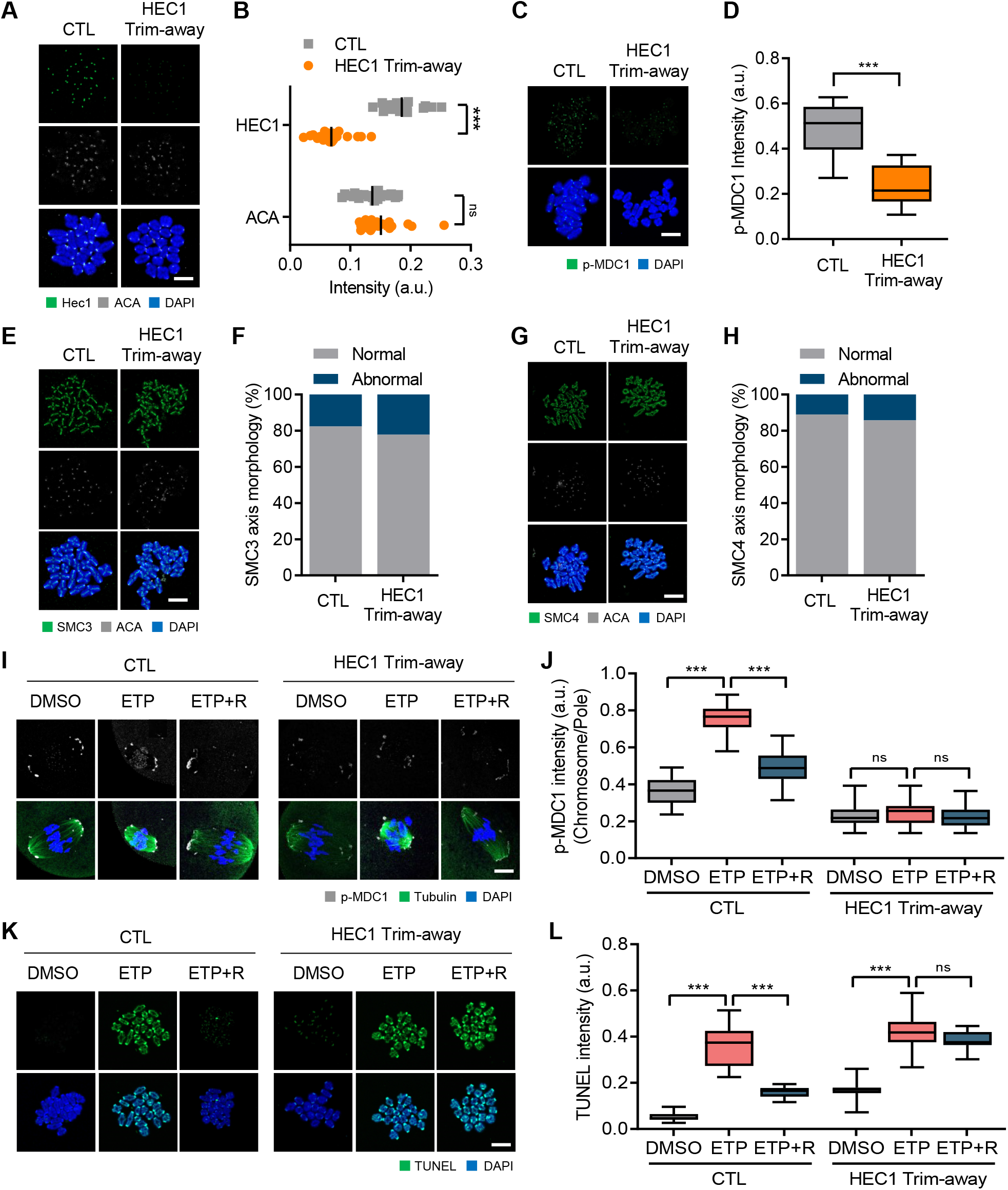
Impaired relocation of CIP2A-MDC1-TOPBP1 complex after DNA damage. (A, C, E, G) Representative images of chromosome spreads from control and HEC1 Trim-away oocytes. Scale bar, 10 μm. Chromosome spreads were stained with HEC1, p-MDC1, SMC3, and SMC4 antibodies. (B, D) Quantification of HEC1, ACA and p-MDC1 intensities. Data are presented as the mean ± SEM from three independent experiments. ***p < 0.0001, ns, not significant. (F, H) Quantification of SMC3 and SMC4 axis morphology. (I) Representative images of control and HEC1 Trim-away oocytes stained with p-MDC1 and tubulin antibodies after ETP treatment and recovery (ETP+R). Scale bar, 10 μm. (J) Ratio of p-MDC1 intensities at chromosomes over spindle poles. Data are presented as the mean ± SEM from three independent experiments. ***p < 0.0001, ns, not significant. (K) Representative images of chromosome spreads showing TUNEL signals from control and HEC1 Trim-away oocytes after ETP treatment and recovery (ETP+R). Scale bar, 10 μm. (L) Quantification of TUNEL intensity. Data are presented as the mean ± SEM from three independent experiments. ***p < 0.0001, ns, not significant.

**Fig. S6.**
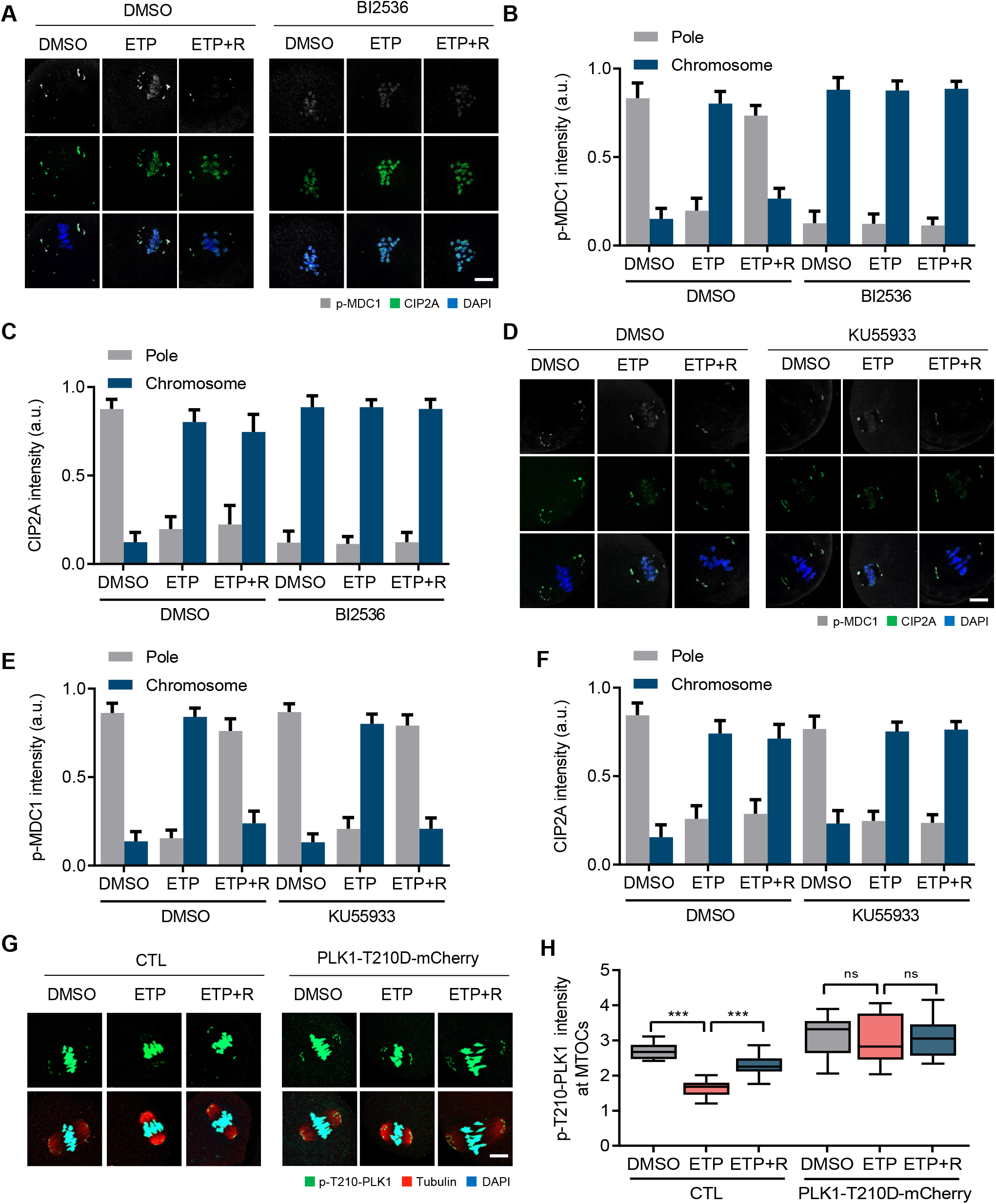
PLK1-dependent relocation of CIP2A-MDC1-TOPBP1 complex. (A, D) Representative images of oocytes treated either BI2536 or KU55933 after ETP treatment and recovery (ETP+R). Control oocytes were treated with DMSO. CIP2A and p-MDC1 signals are shown. Scale bar, 10 μm. (B, C, E, F) Quantification of p-MDC1 and CIP2A localization (chromosome vs pole). Data are presented as the mean ± SEM from three independent experiments. ***p<0.0001, ns, not significant. (G) Representative images showing p-MDC1 localization in oocytes expressing either mCherry or PLK1-T210D-mCherry after ETP treatment and recovery (ETP+R). Scale bar, 10 μm. (H) Quantification of p-T210-PLK1 intensity at MTOCs. Data are presented as the mean ± SEM from three independent experiments. ***p<0.0001, ns, not significant.

**Fig. S7.**
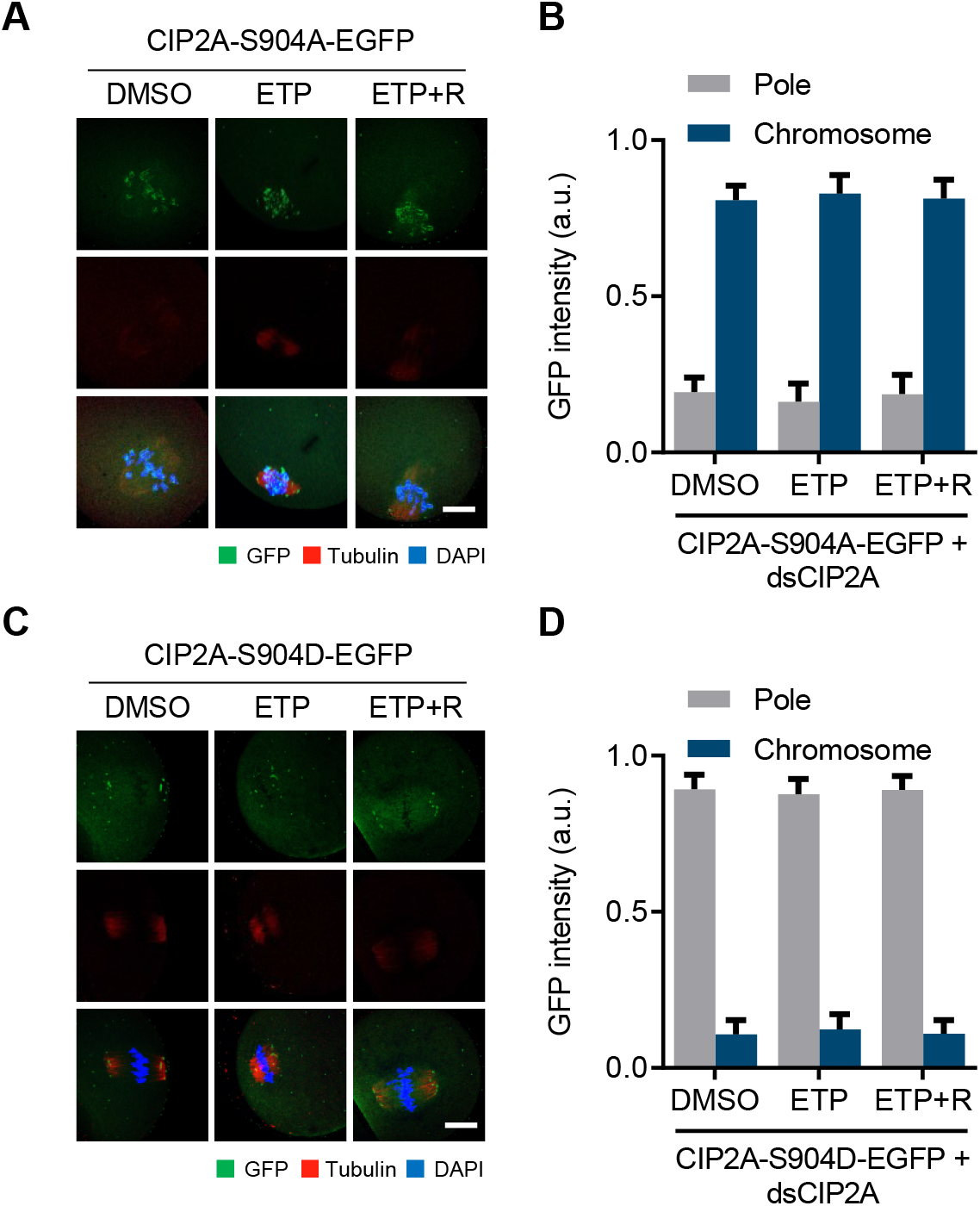
Localization of CIP2A-S904A-EGFP and CIP2A-S904D-EGFP. (A, C) Representative images of oocytes overexpressing CIP2A-S904A-EGFP or CIP2A-S904D-EGFP after ETP treatment and recovery (ETP+R). MI oocytes were stained with tubulin and GFP antibodies. Scale bar, 10 μm. (B, D) Quantification of CIP2A localization (chromosomes vs pole).

